# Genome Dynamics and Chromosome Structural Variations in *Histoplasma ohiense*, a fungal pathogen of humans

**DOI:** 10.1101/2025.05.05.652209

**Authors:** Sarah Heater, Mark Voorhies, Anita Sil

## Abstract

*Histoplasma* is a clinically important but understudied genus of thermally dimorphic human fungal pathogens. *Histoplasma* species normally transition between a multicellular sporulating hyphal form in the soil and a unicellular pathogenic yeast form in a mammalian host. Little is known about genome plasticity of *Histoplasma*, which we address in this study with the ultimate goal of increasing our understanding of its pathogenicity. This study addresses the *Histoplasma* genome and its plasticity to further our understanding *Histoplasma*’s ability to cause severe disease. Here we present the first telomere-to-telomere genome assemblies for *Histoplasma ohiense*. To analyze genome alterations between isolates, we develop an analysis tool to identify genome discontinuities relative to a reference genome. We utilize this analysis to interrogate the genome of laboratory strains and natural *Histoplasma* isolates and discover that the previously published reference genome does not completely match the chromosome structure of the majority of isolates, instead harboring reciprocal chromosome translocations. We identify the telomere-to-telomere *Histoplasma ohiense* reference genome that is most representative of clinical isolates. Additionally, to determine the rate of *Histoplasma* genomic changes, we sequence 46 passaged isolates and calculate the mutation rate to be 2.6 x10^-10^ SNP/base/doubling—the first such measurement to our knowledge within the order Onygenales, which encompasses several critical fungal pathogens. Finally, we sequence populations of cells to assess genome stability over the course of a month under both yeast and hyphal growth conditions. Interestingly, we observe that transposon signal is not static over time and instead increases during growth in both the yeast and hyphal forms as well as through morphologic transitions, suggesting an increase in transposon copy number. Taken together, this work highlights the plasticity of the *Histoplasma* genome and presents a comprehensive genome assembly that is representative of *Histoplasma ohiense* natural isolates.

## Introduction

The order Onygenales contains multiple understudied human fungal pathogens, several of which, including *Histoplasma spp*., rank as priorities on the WHO fungal pathogens list^1^. Species in the genus *Histoplasma* are found globally, including endemic regions of high incidence in Africa and the Americas, with lifetime exposure rates thought to be over 80% in certain hyperendemic regions^1–4^. Although *Histoplasma* can cause life-threatening infections even in healthy adults, severe Histoplasmosis infections are particularly common among immunocompromised individuals with mortality estimates ranging up to 53% in HIV patients with disseminated histoplasmosis^1,5–7^. Treatment options for *Histoplasma* and other Onygenales are insufficient, and vaccines are not yet available^1,5–7^.

*Histoplasma* infections occur when fungal cells are inhaled from the environment, where *Histoplasma* grows in a sporulating hyphal form. *Histoplasma* is thermally dimorphic and, upon inhalation, it has the remarkable capacity to transition to growth as yeast in response to mammalian body temperature^7^. Thus, *Histoplasma* has two ecological niches (soil and mammal) and two temperature-regulated morphological programs (hyphae and yeast). The effect of these morphologies and morphologic transitions on *Histoplasma* genome integrity has not been previously examined.

While *Histoplasma* has been found on all continents, the global incidence of *Histoplasma* is unknown^1,3^. Phylogenetic relationships from sequencing a set of DNA regions suggest at least eight distinct clades of *Histoplasma* globally^8,9^. A few recent studies have performed whole genome sequencing of *Histoplasma* clinical isolates primarily from North America, enabling a more detailed inference of population structure and sequence diversity in this region^10–12^. Based on this sequencing, a handful of *Histoplasma* species (*H. capsulatum* sensu lato) have been proposed^11^. In this study, we will refer to *Histoplasma mississippiense* (also known as NAm1, reference genome derived from clinical isolate WU24) and *Histoplasma ohiense* (NAm2, reference genome from isolate G217B), as names for the two proposed species for which the largest amount of whole genome sequencing data is currently available.

We recently published the first five Nanopore based *Histoplasma* genome assemblies. In this study, we find that our published *H. ohiense* reference genome contains structural differences from most *H. ohiense* clinical isolates and laboratory strains. We develop a more representative *H. ohiense* reference genome and in doing so find indications that *Histoplasma* may have a surprisingly high genome rearrangement rate. Additionally, we assess multiple additional aspects of genome dynamics throughout growth, including mutation rate. Mutation rate is particularly important for pathogens in the context of acquired treatment resistance, and we here make the first measurement of mutation rate in the order Onygenales and the second to our knowledge among Pezizomycotina^13^. We also uncover interesting trends in the copy number of retrotransposons throughout *Histoplasma* growth and morphological transitions. We use these quantified rates of genome plasticity to interpret the observed differences among assembled *Histoplasma ohiense* genomes and to engage in a broader discussion of *Histoplasma* genome plasticity.

## Results

### Analysis of genomic rearrangements in *H. ohiense*

From early studies of *Histoplasma* and recently published genome assemblies, it appears that chromosome size varies among clinical *Histoplasma* isolates from different species^14–16^. However, investigations of *Histoplasma* genome stability led us to the possibility that even very closely related *H. ohiense* strains might also have differences in chromosome composition. By mapping reads from lab strains onto the first Nanopore based *H. ohiense* assembly^16^, we manually identified a point of discontinuity in read coverage on chromosome 2 of the reference assembly compared to all other tested lab isolates. Specifically, we originally identified this point (termed point D) because of a change in read coverage in some isolates (*i.e.,* a copy number variant boundary) and then realized that mapped reads did not span this point. Reads from most strains mapped partially to point D on chromosome 2 and partially to chromosome 6, suggesting that these regions were contiguous in many strains but not in the reference genome. These data suggested a chromosome rearrangement between sequenced strains and the closely related strain used to generate the reference genome. Since understanding the chromosome context is important for genetic manipulations, studies of population structure and genome stability, as well as for various fungal phenotypes, we further investigated this potential rearrangement.

We developed a Jaccard index-based analysis of chromosome continuity of a strain relative to a point along a reference genome (Sequence Continuity Analysis of Reads, SCAR). This analysis was designed to identify points in the genome at which a strain has a structural difference from the reference genome and we applied it to previously sequenced clinical isolates from the literature as well as to laboratory strains. If sequencing reads from a specific strain map evenly around a point on the genome but there is an anomalous gap in reads mapping through this point, SCAR goes to zero, indicating that this strain differs notably from the reference at this point (Fig. 1A). Rearrangements, insertions, or deletions would all cause this type of signal change. If no reads map within 1kb of a tested point (*i.e.,* if the denominator would be zero), SCAR is set to negative, indicating a continuous large deletion. This method is inspired by the Jaccard similarity coefficient in the Trinity program for transcriptome assembly from RNA-seq data. Trinity has a jaccard_clip option for minimizing fusion transcripts derived from gene dense genomes; paired-end reads are mapped to transcript contigs from the first (inchworm) assembly stage and the contigs are clipped at positions spanned by few read pairs relative to the total read pairs proximal to the position^17^. Specifically, the Jaccard similarity coefficient in the Trinity program is defined for a given window as the number of read pairs overlapping both the left window boundary and the right window boundary (*i.e.*, read pairs spanning the window) divided by the number of read pairs overlapping the left boundary, the right boundary, or both boundaries.

**Figure 1.**
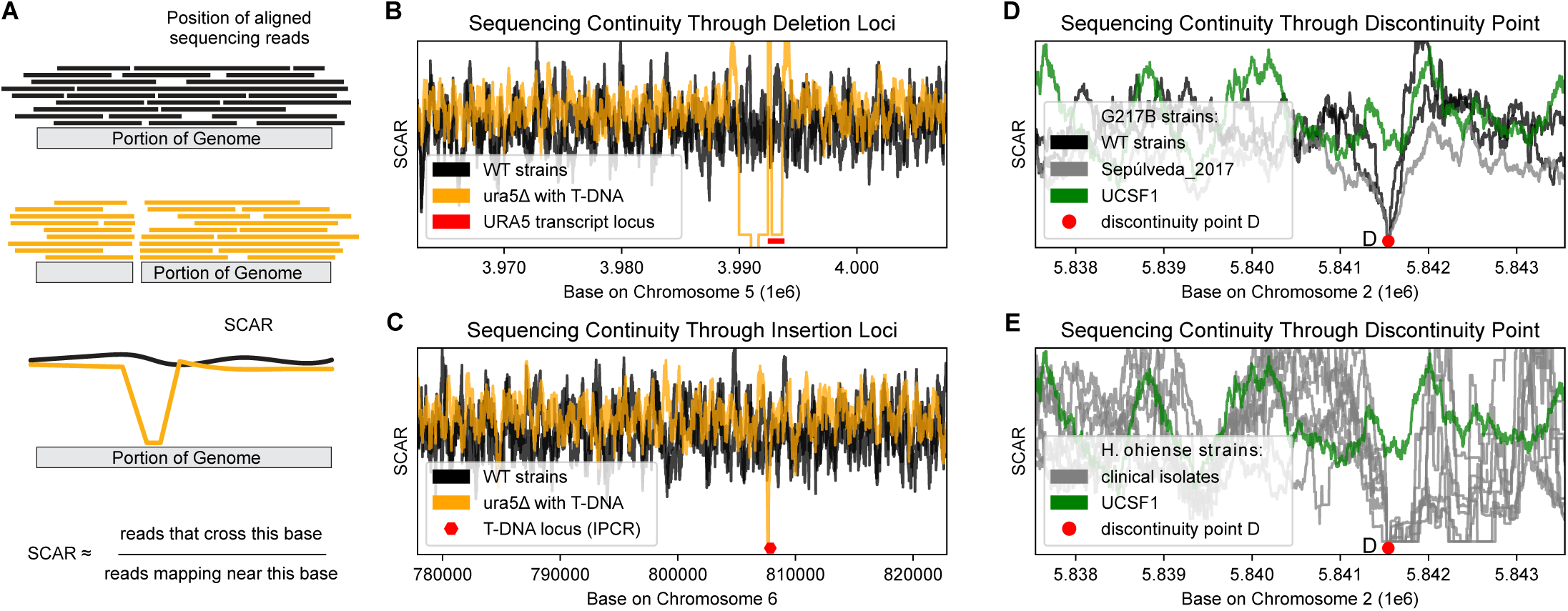
SCAR analysis reveals consistent discontinuity among *Histoplasma ohiense* isolates with respect to the reference genome assembly. A. Schematic of the SCAR analysis used to identify points in the genome at which there is a discontinuity in the position of aligned sequencing reads. B and C. Continuity analysis for WU15-Agr-YB, an example strain harboring an insertion and a deletion. Agr-YB is shown in orange and wild type (WT) strains are shown in black. B. SCAR around the *URA5* locus (red) which is deleted in WU15-Agr-YB. C. SCAR around the WU15-Agr-YB T-DNA insert locus as identified by inverse PCR (IPCR) (shown in red). D) SCAR analysis around point D, shown in red, for representative laboratory strains derived from clinical isolate G217B. E) SCAR analysis around point D for representative clinical isolates. All SCAR assessments in Figure 1 are shown against assembly UCSF1 and, for D and E, the strain from which the UCSF1 assembly was generated is shown in green.

As proof of principle that SCAR can be used to quickly test strains for genomic alterations using available short read sequencing, we applied it to WU15-Agr-YB, a strain with a previously described *URA5* locus deletion^18^. In addition to correctly identifying the deletion (Fig. 1B), this tool was able to pinpoint the locus of an *Agrobacterium* T-DNA insertion which is also present in WU15-Agr-YB (Fig. 1C). Using SCAR, we then confirmed the discontinuity at point D on chromosome 2 that we identified manually, showing that point D is discontiguous for all clinical isolates and for all tested lab strains other than the strain from which the reference assembly was generated (Fig. 1D and 1E). We therefore hypothesized that the strain from which the previous assembly was generated did not have a representative chromosome structure compared to most *Histoplasma ohiense* isolates. Given this finding, we name the previous assembly of G217B “UCSF1” to clearly distinguish it from subsequent assemblies.

### Creation of a new reference sequence for *H. ohiense* and analysis of translocations

To generate an updated reference assembly for *H. ohiense*, we subjected two additional strains to Oxford Nanopore Technologies (ONT) sequencing as well as Illumina sequencing to generate the second and third *Histoplasma ohiense* genome assemblies, UCSF2 and UCSF3. All three strains used to generate *Histoplasma ohiense* reference genomes were derived from the same initial clinical isolate, G217B, and are only separated by minimal laboratory passage. In a first for *Histoplasma ohiense*, we were able to assemble complete telomere-to-telomere assemblies from both newly sequenced strains^16^. As expected, these assemblies are highly syntenic (Fig. 2A, S1, and S2). However, each *H. ohiense* assembly exhibits a slightly different chromosome structure. The region syntenic to the right-hand portion of chromosome 2 of UCSF1 (after point D) joins with a small telomeric section of chromosome 6 to form a separate chromosome in both new assemblies (Fig. 2A). We also detected an additional rearrangement in one of the new assemblies, UCSF3, relative to both other assemblies. As expected, Illumina reads from the strains used to generate both new assemblies did not have continuity by SCAR through the previously described discontinuity point, point D (Fig. 2B) when aligned to genome assembly UCSF1. To address which of these assemblies was most representative of Illumina sequenced natural isolates (used here to represent the broader *H ohiense* population) we also deployed SCAR to assess these natural isolates. All tested clinical and environmental isolates of *H. ohiense* did not have continuity through the UCSF arrangement of point D, termed D-E. These natural isolates instead showed continuity through the alternative arrangement of this point, termed D-d, which is found in assemblies UCSF2 and UCSF3. At the other rearrangement point (arrangement A-a or A-B), most natural isolates agreed with the UCSF3 assembly junction, with discontinuity through rearrangement version A-a and continuity through A-B. We therefore concluded that assembly UCSF3 was most representative of the chromosome structure of natural *H. ohiense* isolates at these rearrangement points.

**Figure 2.**
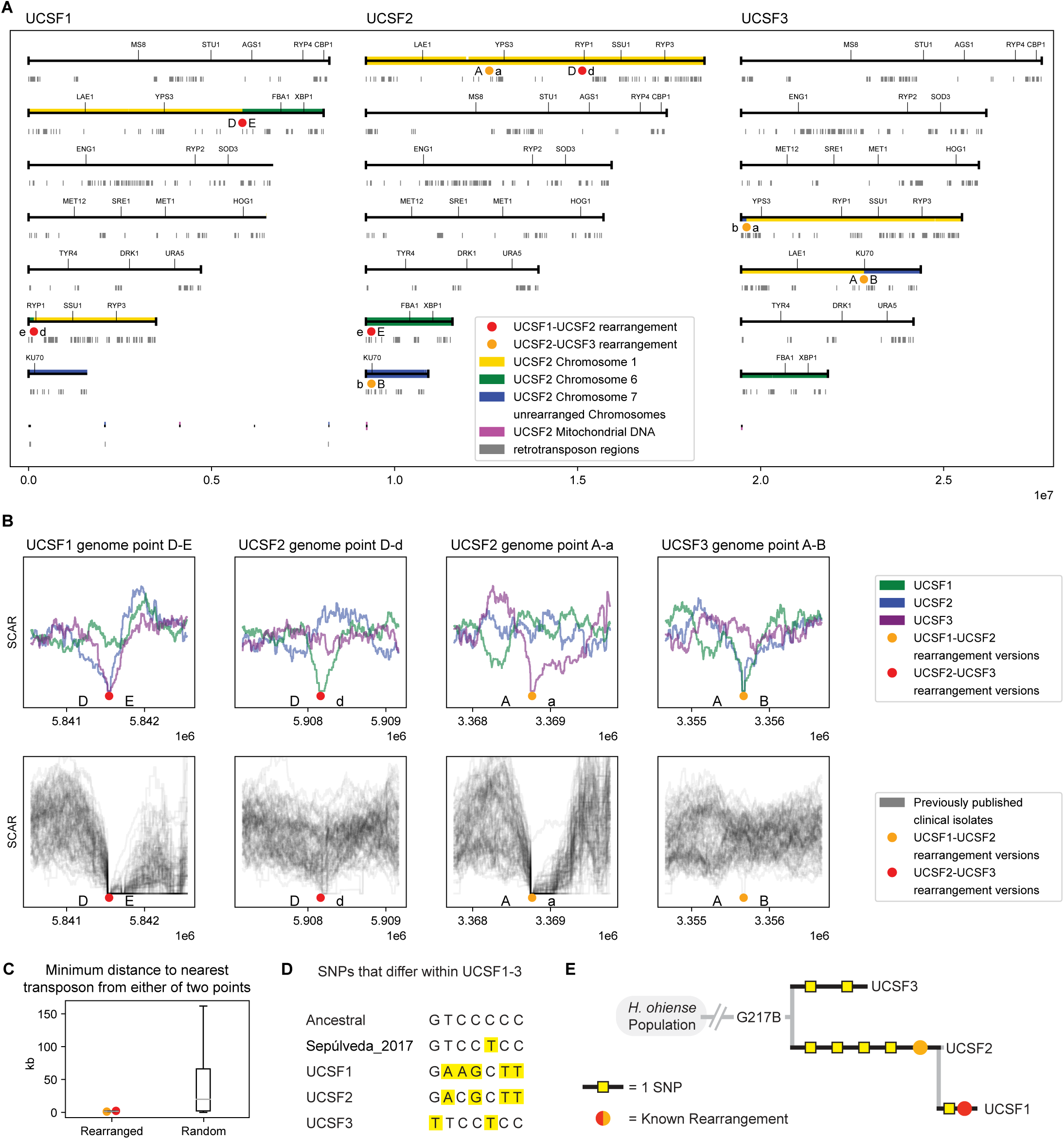
UCSF3 is the assembly most representative of laboratory and natural isolates. Two new genome assemblies were created from alternative isolates of *H. ohiense* G217B. A. Chromosome maps for the three G217B assemblies. Telomeres, if present, are indicated by black vertical lines at the end of each chromosome. The three chromosomes that contain rearrangements between assemblies are highlighted in yellow, green, and blue. NUCMER coordinates were used to color these chromosomes based on synteny to UCSF2. The mitochondrial DNA is shown in purple. Rearrangement points between UCSF1 and UCSF2 genomes are noted with red dots while the rearrangements between UCSF2 and UCSF3 are noted with orange dots. Genes of interest are displayed on each chromosome as landmarks. LTR transposon regions are shown by grey blocks below the chromosomes. B. SCAR analysis at alternative versions of rearrangement points. The illumina reads of the three strains from which the Nanopore assemblies were created are plotted at the top. Below in grey are SCAR plots for previously sequenced *H. ohiense* natural isolates. C. Distance between rearrangement points and nearest LTR transposon versus other non-LTR-transposon points (10000 random sets of two non-transposon points) with rearrangement points colored as in A and B. D. SNP differences among UCSF1, UCSF2, and UCSF3 were identified. These were then compared with the bases at these loci found in other previously published natural isolates (used to infer Ancestral) and a previously published sequence of a G217B isolate (Sepúlveda_2017)^11^. E. Maximum parsimony tree based on SNPs, also showing known rearrangements.

Genomic translocations in other fungi often occur near transposable elements^19–22^. We thus assessed the distance from each rearrangement point to the nearest identified LTR transposon. In keeping with the previously observed trends in other species, these translocation events were particularly close to transposon sequences, at 1.1 kb and 2.2 kb away versus a median of 19.6 kb for 10,000 randomly selected pairs of points (Fig. 2C).

To further assess how these strains relate to each other, as well as to other isolates within the *H. ohiense* population, we subjected UCSF1, UCSF2, and UCSF3 to SNP analysis. Mutations that differ between G217B-derived strains are likely to have arisen during laboratory passage and thus are not expected to be present in the environmental population. Throughout the environmental population, the nucleotide base at each of these loci should be consistent, and that base should reflect the content of the ancestral G217B isolate. We thus identified SNPs that differ amongst these three G217B-derived strains. To infer the most likely state of the common G217B ancestor, these SNPs were compared to those found in other natural isolates of *H. ohiense*. SNPs were also compared to a G217B-isolate-derived strain sequenced by a different group^11^ (Fig. 2D). A maximum parsimony tree of this analysis, also showing the likely acquisition of identified genomic rearrangements, is shown in Fig. 2E. The tree inferred by SNP analysis was congruent with the tree inferred based on rearrangements, with UCSF3 showing the fewest SNPs relative to the inferred ancestral sequence. Because this assembly is the one most likely to resemble both the ancestral state of isolate G217B and *H. ohiense* in the wild, as measured by both SNPs and genome rearrangements, we suggest use of assembly UCSF3 as the reference genome of *H. ohiense* in future genomic studies. We used this assembly for the remainder of this study.

We found it surprising that comparison of the three *H. ohiense* assemblies revealed two translocations and 7 SNPs (Fig. 2E). We are not aware of an analysis of the relative rates of SNP and translocations amongst fungi, especially through mitotic growth, but the mutation rate in the model fungi *S. cerevisiae* is thought to be much higher than the rate of genomic translocations during mitotic growth^20^. While we cannot retrospectively determine the number of doublings that separate the strains used to generate these *H. ohiense* assemblies, our findings suggested that during the laboratory passaging separating these isolates, the mutation rate might be unusually low or the translocation rate might be particularly high.

### First quantification of mutation rate in *Histoplasma*

The mutation rate in *Histoplasma* and in other fungi of the order *Onygenales* has not been previously measured. To assess SNPs per doubling, *H. ohiense* was grown up briefly in vitro while cell doublings were counted followed by identification of SNPs in offspring (Fig. 3). Doublings were monitored by counting cells with a hemocytometer as *H. ohiense* yeast was grown up first as single colonies then briefly in liquid culture. Offspring were then grown up from single colonies and subjected to whole-genome Illumina sequencing. The sequences of “sibling” offspring were compared to each other to find mutations in particular offspring versus the offspring-consensus-inferred parental genome. Two parental genomes were also sequenced to spot check accuracy of offspring-inferred parental genomes, confirming that we could correctly infer parental SNP content from concordance among offspring. As is standard for calculations of mutation rate, repetitive genomic regions that were highly enriched for transposons were excluded from mutation rate analyses due to ambiguous Illumina read mapping in these regions^20^. We determined that the mutation rate in *H. ohiense* is 2.6 x10^-10^ SNP/base/doubling (95% CI 1.2x10^-10^ to 4.9x10^-10^). This rate is similar to what is found in model yeast but slightly higher than observed in *Aspergillus*, the most closely related species for which a measured mutation rate is available^13,20^.

**Figure 3.**
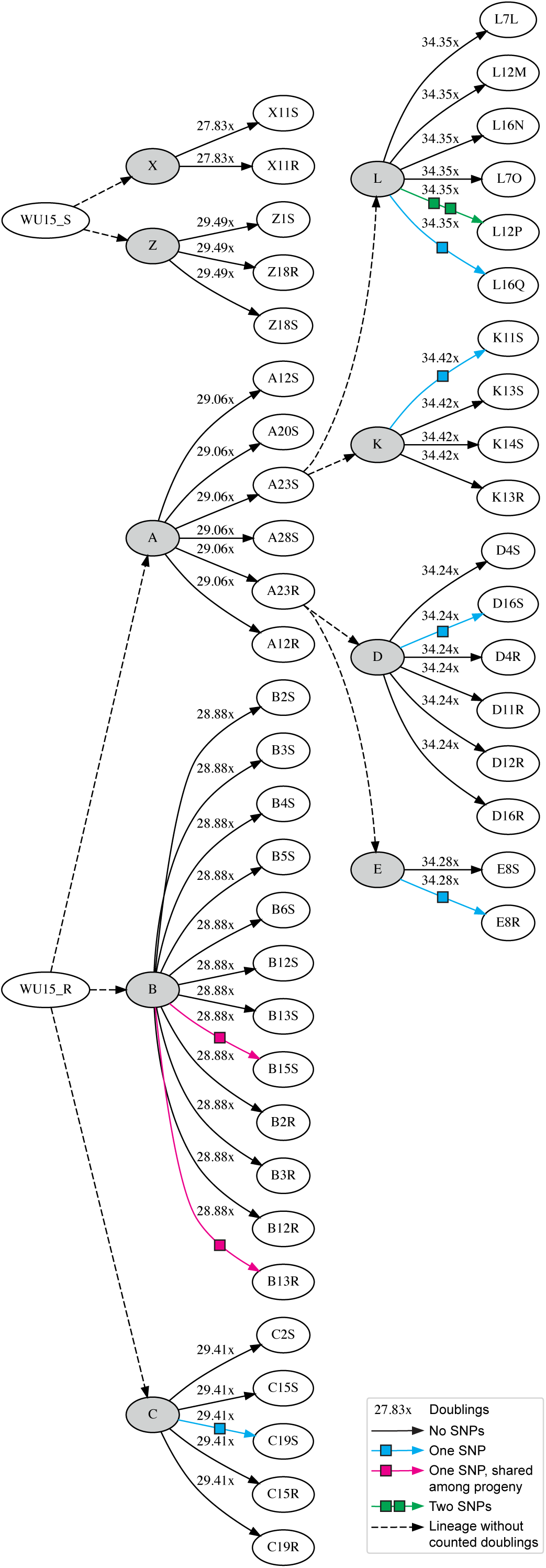
Schematic illustrating pedigree chart used to determine mutation rate. Parental colonies (grey ellipses) were grown up briefly in liquid culture, and hemocytometer counts were used to determine doublings (noted above arrows) from the point of a single parental cell. Offspring cells from this culture were grown up as single colonies then in liquid culture for whole genome sequencing which was used to identify SNPs (squares along arrows). The number of SNPs identified was used in concert with the number of doublings counted to find SNPs per base per doubling.

### Analysis of retrotransposons in *Histoplasma* assemblies, species, and through growth stages

Previous studies have identified transposable elements within *Histoplasma* species, primarily a Ty3/Gypsy long terminal repeat (LTR) retrotransposon^16,23,24^. However, transposon content has not previously been studied through cell growth nor has it been compared *en masse* amongst sequenced clinical isolates of multiple species. Given that temperature can stimulate transposon mobility in *Cryptococcus*^25^, and transcript abundance of transposons and transposon-embedded genes is increased in yeast versus hyphal *Histoplasma*^16^, we asked if LTR transposon proliferation varies with morphological growth state as *Histoplasma* transitions between morphologies. Starting with isolates of wild type *H. ohiense ura5Δ*, we grew *Histoplasma* over the course of 4 weeks as yeast (maintained by growth at 37°C), as hyphae (maintained by growth at 25°C), and through morphology transitions (yeast-to-hyphae and hyphae-to-yeast, induced by temperature shift at the start of transition) to assess the retrotransposon signal. Samples were taken for whole-genome Illumina sequencing every two weeks for steady-state yeast or hyphae and once a week for the morphology transitions. Relative LTR transposon signal was assessed by normalized coverage of Illumina reads aligned to a single full copy of this retrotransposon. Samples were also fixed for DIC microscopy throughout the transitions and scored for morphology to determine the timepoint at which the morphology changed (Fig. 4A and S3).

**Figure 4.**
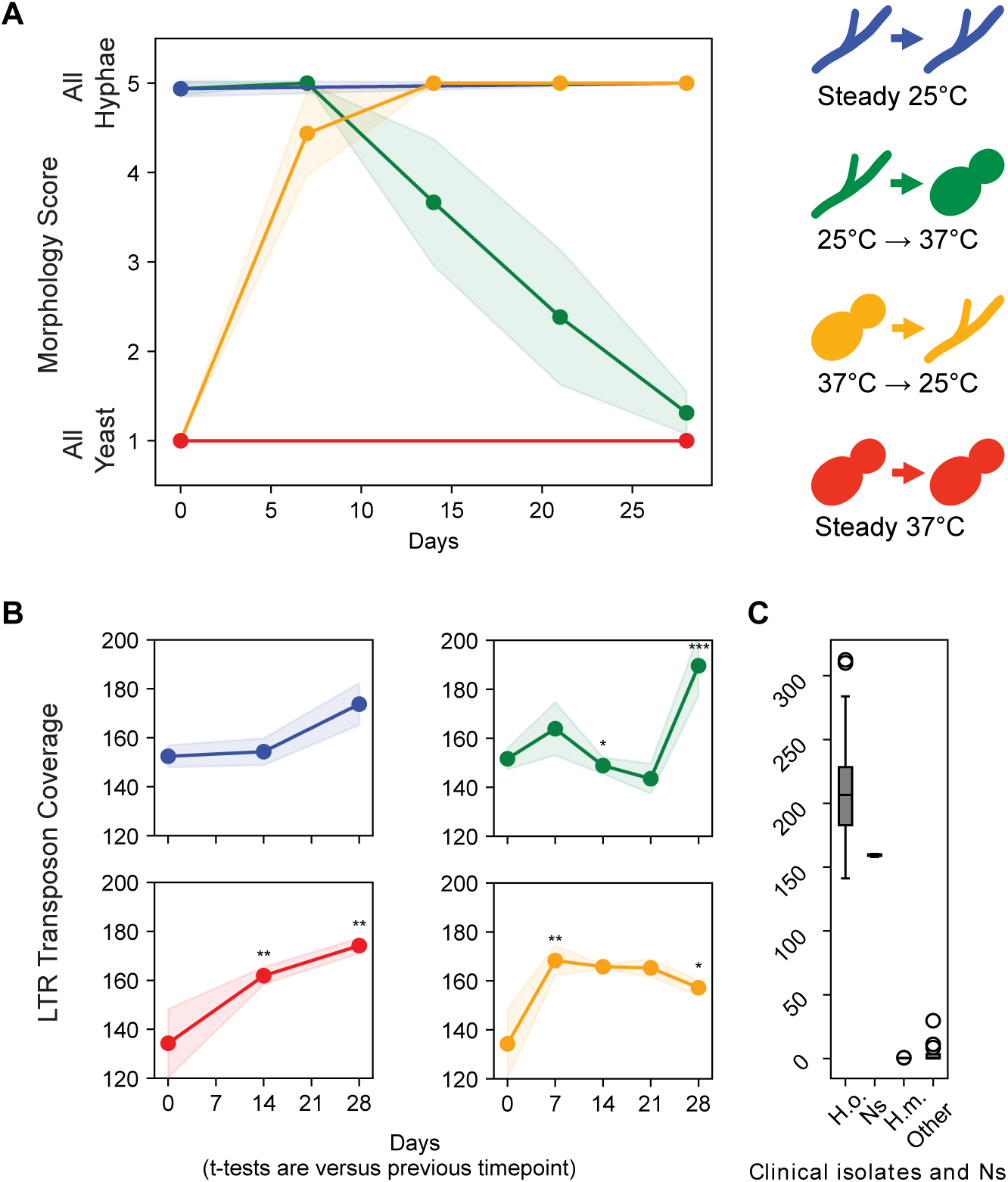
Changes in transposon content during *in vitro* growth including morphological transitions. A. Morphology through in vitro growth of steady-state hyphae (blue), the hyphae to yeast transition (green), the yeast to hyphae transition (yellow), and steady-state yeast (red). Shaded regions indicate standard deviation at each assessed point. B. Normalized coverage of LTR transposon DNA through in vitro growth in each of the four morphology states or transitions, with standard deviation indicated by color shading as in A. Asterisks indicate significance by t-test by comparison to previous timepoint. Clockwise from top left: steady-state hyphae, hyphae to yeast, yeast to hyphae, steady-state yeast. P-value equal to or above 0.05 is not considered significant. P-values < 0.05 are indicated by *, p-values < 0.01 are indicated by **, p-values < 0.001 are indicated by ***. C. Normalized coverage of LTR transposon DNA also shown for natural isolates of *H. ohiense* (*Ho*), the 3 strains from which *H. ohiense* Nanopore-based genomes were created (Ns), natural isolates of *H. mississippiense* (*Hm*), and natural isolates in other categories.

Surprisingly, LTR retrotransposon signal increased throughout all four *in vitro* growth conditions (Fig. 4B). On average over all time courses and conditions, there was a 22% increase in normalized coverage over the course of four weeks. In the transition experiments, a sharp increase in retrotransposon signal correlated with the final conversion to yeast morphology (32% increase from day 21 to 28) as well as with the final conversion to hyphal morphology (25% increase from day 0 to 7). The three Nanopore sequenced G217B isolates had a very similar transposon content, which was close to the median transposon content during the time courses. The transposon content of G217B was in the lower quartile of the transposon content observed in sequenced natural isolates of *H. ohiense*, suggesting that higher transposon content may be typical in the *H. ohiense* natural population. Consistent with trends observed amongst the previous *Histoplasma* genome assemblies, *H. ohiense* isolates universally had higher LTR transposon content than *H. mississippiense* isolates (Fig. 4B)^16^.

## Discussion

The ability to study *Histoplasma* biology is dependent on a representative genome assembly. Our analysis of genome contiguity followed by *de novo* genome assembly of closely related *Histoplasma* isolates revealed that each assembled isolate contained a slightly different set of chromosomes. Chromosomal rearrangements affect the context in which genes lie and can affect gene expression, drive gene evolution, and are associated with fungal characteristics including morphology, fitness, aneuploidy, and antifungal resistance^26^. The initial impetus for our study was the identification of a genomic discontinuity defined by sequencing and analysis of natural and laboratory isolates. We were able to confirm this discontinuity via the generation of new Nanopore-based genome assemblies and developed parameters for an assessment of such discontinuities amongst additional short-read sequenced isolates (SCAR). By comparing our assemblies to genome discontinuities among sequenced natural isolates and laboratory strains, we determined that the chromosome structure of *H. ohiense* UCSF3 was most representative of almost all sequenced strains. We subsequently found that UCSF3 is also most representative at the nucleotide level by SNP analysis.

In addition to being representative of *H. ohiense* isolates, UCSF3 also demonstrates the most synteny with assemblies of other *Histoplasma* species. For example, chromosome 5 of UCSF3 contains both *LAE1* and *KU70*, which are located on separate chromosomes in UCSF1 and UCSF2. A chromosome syntenic to UCSF3 Chr5, also containing both *LAE1* and *KU70*, is found in the assemblies of HcH88, HcWU24, and HcG186AR. Similarly, a chromosome syntenic to the shortest chromosome of UCSF3, Chr7 which contains *FBA1*, is found in the assembly of HcWU24 (*H. mississippiense*) while this region is attached to Chr2 of UCSF1 in a join not observed in other species’ reference genomes. Previously, three regions of notable synteny were described among *Histoplasma* and other closely related pathogens^16^. These three regions are located on three of the four chromosomes not affected by the UCSF1, UCSF2, and UCSF3 rearrangements. One of the clearest regions of synteny among *Histoplasma* strains and species is on Chr3 in UCSF3, which is also conserved out to *Blastomyces*. This region contains most genes involved in sulfur assimilation (including MET1 and MET12) and the mating type locus (MTL), and is not rearranged between UCSF1, UCSF2, and UCSF3. The highly conserved region around DRK1 and TYR1 (on Chr6 of UCSF3) which is also conserved in *Blastomyces*, *Paracoccidioides*, and *Coccidioides* is similarly consistent among UCSF1, UCSF2, and UCSF3. This is also true of the end of UCSF3 Chr1 (i.e. UCSF1 Chr1 and UCSF2 Chr2) which contains *CBP1, HSF1* and *RYP4*. More sequencing is needed to determine if translocations in these regions are significantly less common than translocations elsewhere in the genome.

We also assessed SNP rate and found that the mutation rate in *Histoplasma* is similar to mutation rates previously identified in other fungi and in other kingdoms of life. It remains to be seen if the rate might differ in hyphae or during morphological transitions, which is a technically challenging assessment since hyphal growth is by definition a multicellular form.

In assembling these genomes, we were surprised by the number of translocations observed among such closely related isolates, suggesting that during mitotic laboratory growth, the rate of *Histoplasma* translocations may be particularly high relative to model yeast. It is likely that a multitude of factors could influence translocation rates--for example, strong selective pressure during experimental evolution of fungi is associated with an increased rate of genomic translocation^22^ and it is possible that standard laboratory passage of *Histoplasma* triggers an unknown selection. However, even if the typical translocation rate is high through mitotic growth of *Histoplasma* in nature, cells with certain rearrangements may be selected against during meiosis, limiting the translocations among the environmental population and enabling the observed high degree of synteny between *Histoplasma* species. A potential high mitotic translocation rate in *H. ohiense* could be associated with its high transposon content. Genomic rearrangements in other fungi are often adjacent to transposable elements^20–22^, and *H. ohiense* genomes contain a high transposon content (20%) relative to other sequenced *Histoplasma* isolates and relative to model yeast (3%)^16,27^. We find that rearrangement points between these three assemblies are on average closer to a transposon than an average point within the genome, although the total count is too small for this to reach significance. In each of the identified translocations, one side of each rearrangement is found less than 2.5 kb from a transposon.

Although it has been known for some time that the LTR transposon content of some *Histoplasma* genomes was high, the evolution of transposon content within a lineage has not been previously studied. We observed persistent increases in LTR transposon signal through growth under 4 conditions, with sharp upticks in signal corresponding generally to the timing of visual conversion in morphology (either to hyphae or to yeast). We interpret this increase in signal to reflect an increase in transposon content in each lineage. This work makes clear that growth immediately prior to sequencing can rapidly change the retrotransposon signal which may contribute to the variability in retrotransposon content between previously sequenced isolates. Since LTR transposon content in *H. ohiense* cannot increase infinitely, we hypothesize that there is either a limit to transposon expansion or some biological context where transposon content contracts. While meiosis provides possibilities for transposon content reduction in nature^28^, we have also found that laboratory strains which have been subjected to significant mitotic passaging and manipulation occasionally have a significant contraction in their LTR transposon content as assessed by genome sequencing. The mechanism for this contraction is unknown, but programmed and reproducible massive removal of genome portions including transposable elements has been observed in other species^29,30^. Certain mechanisms limiting transposon proliferation may also be more abundant in nature and in the presence of meiosis including repeat-induced point mutation (RIP) and meiotic silencing^28^. Future studies are needed to understand the parameters that govern *Histoplasma* transposon expansions and contractions.

We posit that the parameters that govern transposon content may be different in *H. ohiense* versus other *Histoplasma* species, which tend to have much lower LTR transposon coverage as indicated by sequencing. It is possible that machinery related to transposon contraction or expansion may differ between these species. Highly divergent retrotransposon content has been previously observed amongst other closely related fungal species, such as a stark difference in LTR retrotransposon content between two species in the genus *Blastomyces* that is hypothesized to play a role in speciation^31^. Transposons have been observed to sometimes confer fitness advantage to their hosts, for example in *Schizosaccharomyces pombe*, depending on growth conditions and mating type^32^. Differential transposon content amongst *Histoplasma* species may be related to different evolutionary pressures or relative advantage conferred by transposons in the context of other genetic and cellular differences, which can be explored in future studies.

**Supplemental Figure 1.**
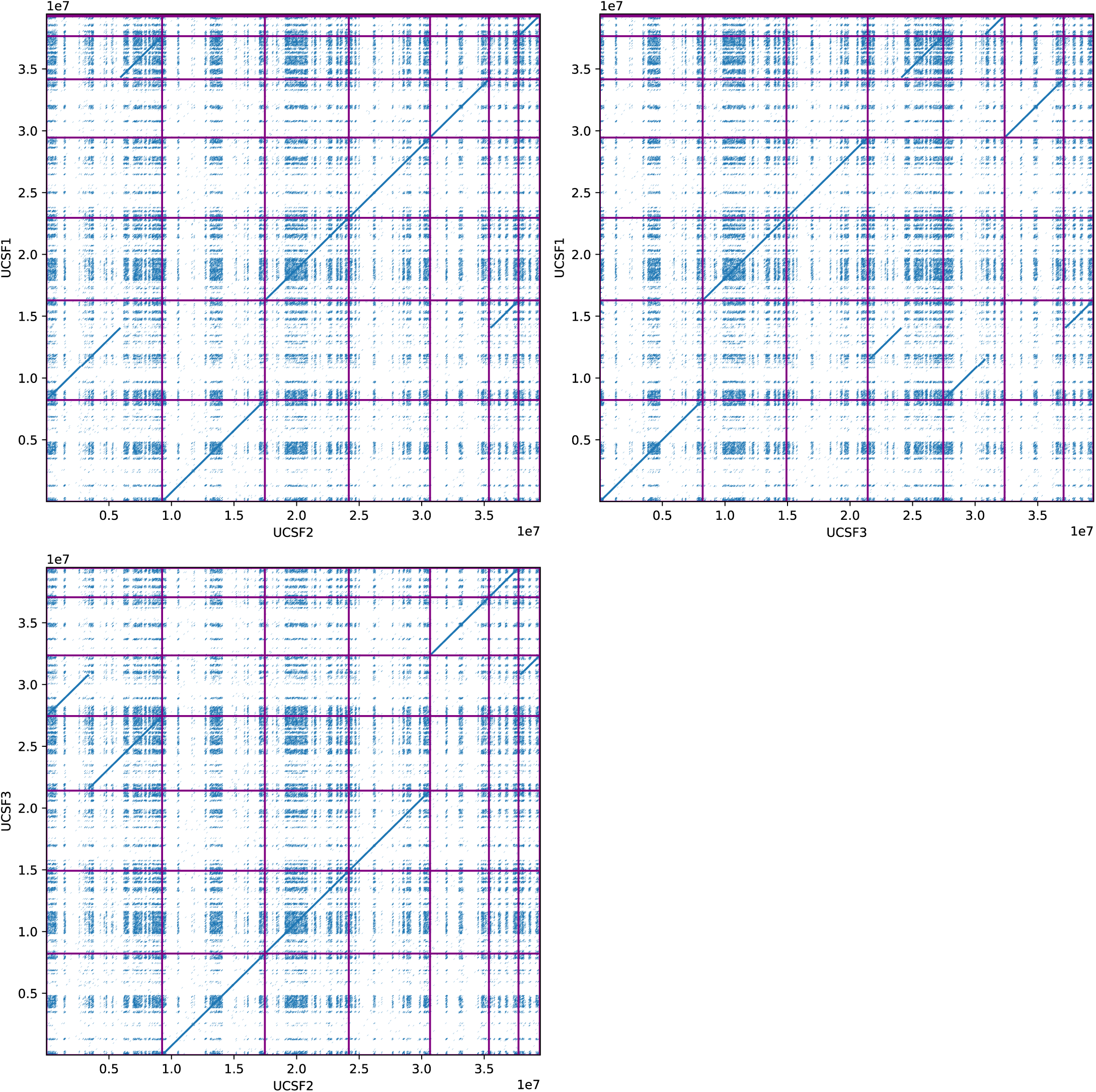
NUCMER Dotplot comparisons of the three *Histoplasma ohiense* genome assemblies

**Supplemental Figure 2.**
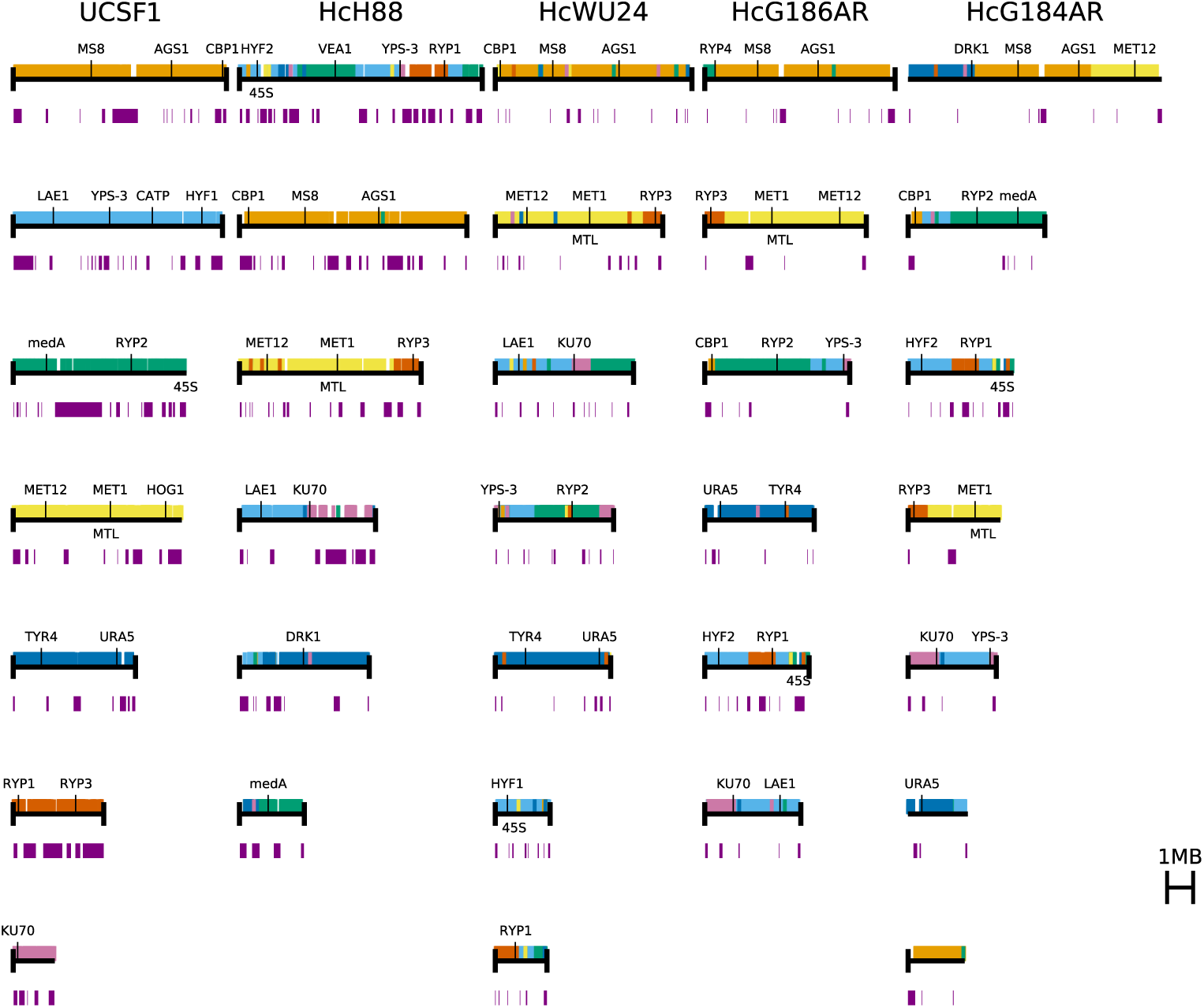
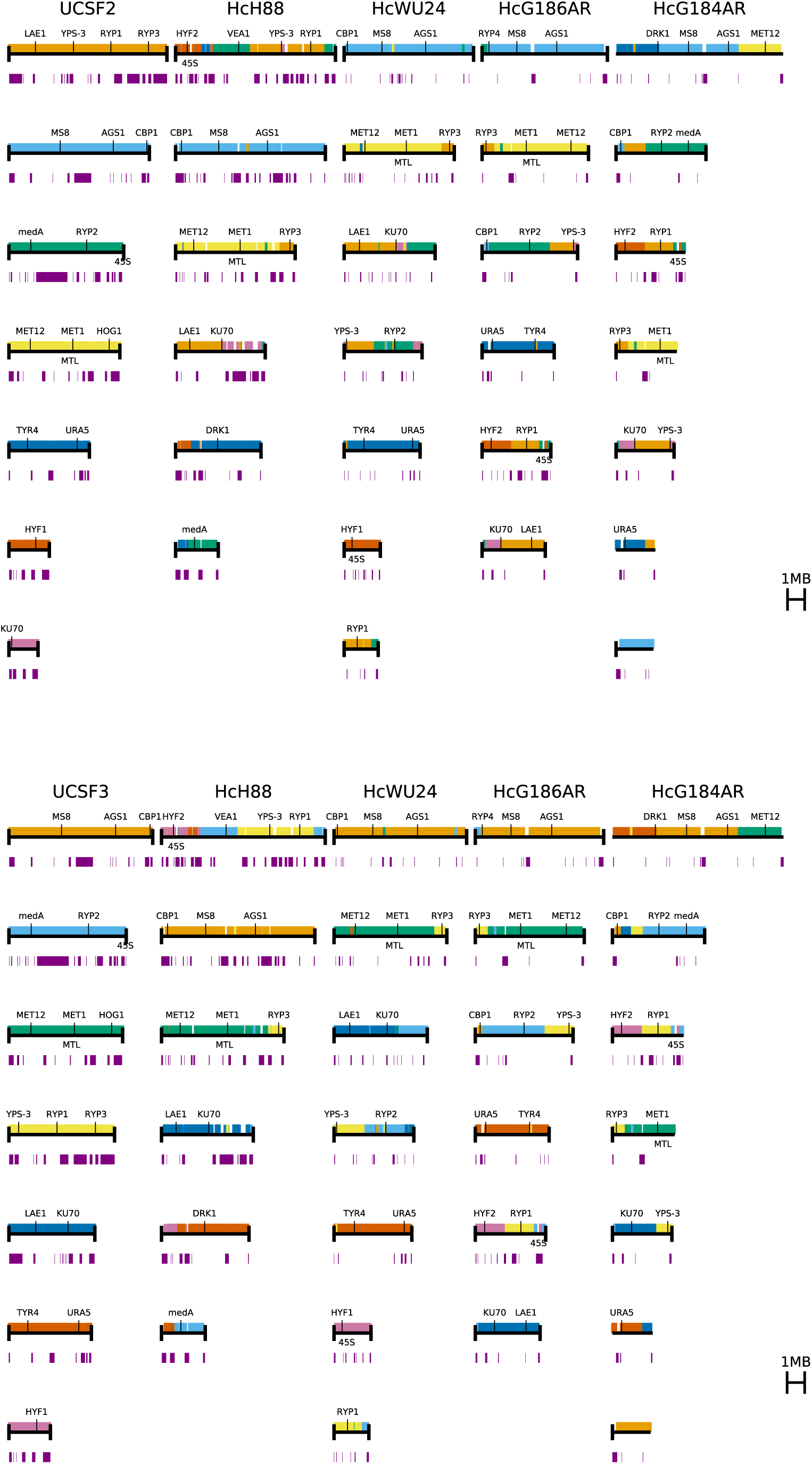
Regions of synteny between UCSF2 or UCSF3 and the *Histoplasma* assemblies from Voorhies, 2022.

**Supplemental Figure 3.**
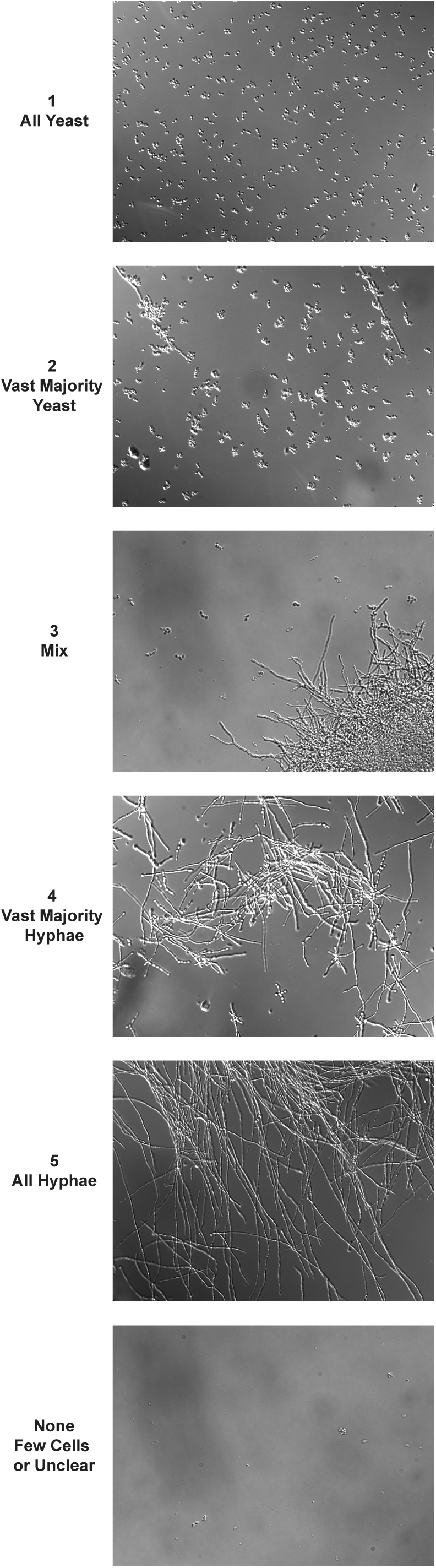
Legend used for scoring of microscopy images

**Supplemental Table 1.** Strains used in this paper

**Supplemental Table 2.** Gene annotations among genome assemblies

**Supplemental Code.** SCAR index

## Materials and Methods

### Histoplasma strains

WU15-Agr-YB (used as a representative strain harboring an insertion and a deletion) was generated using the *Agrobacterium-*based insertional mutagenesis method described in Rodriguez, et al^33^. Inverse PCR (IPCR) was used to estimate the location of the T-DNA insertion prior to confirmation by whole genome sequencing and SCAR analysis. UCSF1 UCSF2 and UCSF3 are each minimally passaged strains of *H. ohiense* clinical isolate G217B. Previously sequenced natural isolates of *Histoplasma* used as reference include isolates defined as *H. ohiense* (or as NAm2), those defined as *H. mississippiense* (or as NAm1) and those defined as belonging to other *Histoplasma* categories amongst three studies that previously published *Histoplasma* clinical isolate sequences: Bagal et al, Tenório et al, and Sepúlveda et al^10–12^. Strains used to define mutation rate and for analysis of variable copy number regions were from the *Histoplasma* G217B *ura5Δ* (WU15) background. To assess transposon content, strains with and without copy number alteration were assessed as follows: For timecourses of yeast, hyphae, and through morphology transitions, a 1:1 mix of strains with and without a copy number alteration were used. This copy number alteration is the subject of a manuscript currently in preparation. Strains used in this paper can be found in supplemental table 1.

### *Histoplasma* general growth conditions, DNA extraction, and sequencing

*Histoplasma* growth conditions were as described previously unless otherwise indicated^34^. Extraction of genomic DNA for Illumina short read sequencing was performed as described previously for yeast samples^34^. Hyphal samples and samples from throughout morphological transitions were collected under biosafety level 3 conditions utilizing a slightly modified DNA extraction protocol. For these samples, cells were collected by filtration^34^ rather than by centrifugation and cells were bead-beat 2-3 times rather than once, or until the mixture appeared smooth. For all DNA extractions for short read sequencing, in any instance where a visible DNA precipitate did not appear after addition of isopropanol, to ensure ample DNA was extracted, 1.5 μl GlycoBlue Coprecipitant (Invitrogen™ AM9515) was added followed by an overnight incubation at 4 °C. Subsequent centrifugation steps were done at 4 °C instead of at ambient temperature. Illumina library preparation and sequencing was either performed using a Nextera DNA Flex Library Prep kit followed by sequencing at the UCSF Center for Advanced Technology or performed by SeqCenter, LLC (Pittsburgh, PA).

For extraction of genomic DNA to be used for long read Nanopore sequencing and associated short-read sequencing of these strains, 2-10 mL of a 3 day liquid culture was pelleted (2500 rpm 5 minutes), washed once in TE buffer (spin at 13000 rpm for 1 minute), then stored at -80 °C until DNA extraction. DNA extraction was then performed using components of the Qiagen Gentra Puregene Yeast/Bacteria kit (158567) with a slightly modified protocol. Wide-mouth pipette tips were used when pipetting cell lysate or DNA, and vortexing was avoided other than as noted. *Histoplasma* cell pellets were resuspended in 300 μl Cell Suspension buffer, approximately 100 μl 0.5 mm glass beads were added, the mixture was thrice vortexed for 10 seconds and put on ice for 10 seconds between each vortex. 4 μl of the Lytic enzyme was then added, this mix was inverted 25 times, then incubated for 2 hours at 37 °C with inversion 3 times an hour. This mixture was then pelleted at max speed for 1 minute at 4 °C, supernatant was discarded, 300 μl Cell Lysis Solution was added, then the mix was slowly pipetted several times and incubated at ambient temperature for 10-20 minutes. 100 μl Protein Precipitation Solution was then added, the mixture was immediately vortexed for 5 seconds, placed on ice, vortexed for 1 second, then placed on ice for 5 minutes. This mix was pelleted at max speed for 3 minutes at 4 °C. Supernatant was then transferred to a tube containing 300 μl isopropanol and inverted 50 times to mix. This sample was pelleted at max speed for 2 minutes at 4 °C. Supernatant was removed leaving approximately 50 μl so as not to disturb the DNA pellet. 500 μl 70% Ethanol wash was added, then DNA was pelleted at max speed for 1 minute. Supernatant was fully removed, pellet was air dried for 15 minutes, then 70 μl DNA Hydration Solution was added along with 1.5 μl RNase A Solution. This was gently mixed by slowly pipetting up and down 10 times, then incubated at 37 °C for 5 minutes, 65 °C for 10 minutes, then overnight at 4 °C to dissolve the pellet. Nanodrop and TapeStation (Agilent Genomic DNA reagents) were used to assess quality and concentration prior to library preparation and sequencing at SeqCenter, LLC.

### SeqCenter sequencing

Illumina sequencing libraries were prepared using the tagmentation-based and PCR-based Illumina DNA Prep kit and custom IDT 10 bp unique dual indices (UDI) with a target insert size of 280 bp. No additional DNA fragmentation or size selection steps were performed. Illumina sequencing was performed on an Illumina NovaSeq X Plus sequencer in one or more multiplexed shared-flow-cell runs, producing two 151 bp paired-end reads. Demultiplexing, quality control and adapter trimming was performed with bcl-convert1 (v4.2.4).

Nanopore sample libraries were prepared using the PCR-free Oxford Nanopore Technologies (ONT) Ligation Sequencing Kit (SQK-NBD114.24) with the NEBNext® Companion Module (E7180L) to manufacturer’s specifications. No additional DNA fragmentation or size selection was performed. Nanopore sequencing was performed on an Oxford Nanopore GridION sequencer using R10.4.1 flow cells in one or more multiplexed shared-flow-cell runs. Run design utilized the 400 bps sequencing mode with a minimum read length of 200 bp. Adaptive sampling was not enabled. Guppy 1 (v6.5.7) was used for super-accurate basecalling (SUP), demultiplexing, and adapter removal (dna_r10.4.1_e8.2_400bps_modbases_5mc_cg_sup.cfg).

### Genome assembly and annotation

For each isolate (UCSF2 or UCSF3), reads were independently assembled by FLYE version 2.9.4-b1799 with parameters --asm-coverage 40 --genome-size 40m –nano-hq flags and CANU v2.3-development (git commit c61ebbb7) with parameters genomeSize=40m and -nanopore. For both isolates, the FLYE and CANU assemblies were highly congruent, as assessed by NUCMER alignments (using NUCMER from MUMMER 3.23). The CANU assemblies were more complete with respect to telomeres and were used as the basis for manual finishing.

The UCSF2 CANU assembly had 7 large contigs with 6 having telomeres on both ends and one (contig 4) having a telomere on only one end. A small contig with the final UCSF2 telomere was identified by NUCMER alignment of the UCSF2 CANU assembly to the UCSF3 FLYE assembly, and this small contig was appended to UCSF2 contig 4 based on the overlapping bases at the ends of the two contigs.

The UCSF3 CANU assembly had 9 large contigs with 3 having telomeres on both ends and 5 (contigs 2, 3, 6, 8, and 9) having a telomere on only one end. A join between contigs 6 and 8 was identified by NUCMER alignment of the UCSF3 FLYE and CANU assemblies. A join between contigs 3 and 9 was identified by NUCMER alignment of the UCSF2 and UCSF3 CANU assemblies. In both cases, joins were applied by concatenating the incomplete contigs on overlapping bases. Finally, we observed that contig 2 of the CANU assembly of UCSF3 ended in the 45S rDNA repeat whereas the syntenic region in the UCSF2 assembly ended in the 45S rDNA repeat immediately followed by a telomere repeat with no intervening sequence. Therefore, we concatenated a telomere repeat to the 45S rDNA repeat of UCSF3 contig 2. After PILON polishing with UCSF3 Illumina reads (see method below), we validated this extension by aligning UCSF3 Nanopore reads to the PILON polished assembly. Indeed, this identified two reads longer than 50kb that include the 45S rDNA repeat followed by a telomere repeat with no intervening sequence.

For both isolates, CANU assembled many copies of the mitochondrial DNA (mtDNA) as concatamers of different copy number. These mtDNA contigs were identified by alignment to the known mtDNA of the Sanger assembly of *H. capsulatum* G186AR (GCF_000150115.1 supercont1.87). For each isolate, a single mtDNA contig was selected and was clipped to single copy and circularized based on self comparison with NUCMER. For both isolates, after manual finishing of telomere-to-telomere nuclear chromosomes and the mtDNA, the remaining CANU contigs contained no unique sequence and were discarded.

Strand assignments were chosen to put syntenic chromosomes from UCSF2 and UCSF3 in matching orientations and the assemblies were polished with paired end Illumina reads from the corresponding isolate by up to 10 iterations of BWA MEM version 0.7.17 and PILON version 1.23. Finally, remaining Nanopore adapter sequence was identified and removed by trimming each nuclear chromosome to its terminal telomere repeats.

Final assemblies were annotated as in Voorhies, 2022^16^: rRNA was annotated with RNAMMER 1.2 in eukaryote mode, tRNA was annotated with TRNASCANSE 2.0.7, LTR transposons were annotated with LTRHARVEST from GENOMETOOLS 1.6.1 and TBLASTN from NCBI BLAST+ 2.11, and mRNAs were mapped from Gilmore, 2015^35^ using BLAT v35 followed by repair of disrupted reading frames using TBLASTN and MEGABLAST from NCBI BLAST+ 2.11. As before transposon-rich regions, which we refer to as LTR blocks, were defined by joining transposons within 50kb of each other^16^.

### Sequencing continuity analysis by SCAR

See supplemental code. In short, this is defined for each point in a sequenced sample aligned against a reference genome as the number of read pairs where alignment region overlaps this point divided by read pairs where alignment region contains at least one base within 1 kb of the tested point. The 20 bp on each end of an aligned read set are excluded to allow the index to consistently go fully to zero even in cases where short similarities are present on either side of the rearrangement. Read pairs where coverage region is vastly outside of the standard range (over 100 kb or under 50 bp) are likewise excluded from the analysis. If no read pair coverages are found within 1 kb of a point (i.e. the numerator and denominator would both be 0) output is instead set to -0.1.

### Visualization of syntenic regions among assemblies

Pairwise sequence alignments among the three G217B assemblies were generated using NUCMER with default parameters and alignment coordinates were extracted from the resulting delta files using show-coords -r -c -l. These alignments are plotted in Fig. S1 as lines joining the start and end coordinates in each genome. For Fig. 2A, the coordinates of the 1000 longest alignments to UCSF2 chromosomes 1, 6, and 7 were plotted relative to each genome and colored by UCSF2 chromosome. In cases where a genomic region was included in multiple alignments, the longest alignment is shown. For the synteny visualization among distinct *Histoplasma* species in Fig. S2, colors were assigned by chromosome for a given reference genome (UCSF2 or UCSF3) and orthologous genes in each genome were colored by these assignments, as in Voorhies *et al* 2022^16^.

### Analysis of retrotransposon locations and of distance to nearest retrotransposon region

For distance analysis for each translocation, the nearest LTR transposon locus was found on either side of the original pre-translocation points in the UCSF3 assembly. For comparison, the nearest repeat locus was also determined for each of 10,000 random sets of two non-repeat loci.

### Inference of UCSF1/UCSF2/UCSF3 lineages relative to G217B

Paired-end Illumina reads from UCSF1, UCSF2, UCSF3 and 93 non-G217B *H. ohiense* population isolates were aligned to the UCSF2 reference using BWA MEM 0.7.17 with default parameters. A SNP matrix was defined from the UCSF1, UCSF2, and UCSF3 read alignments as nucleotide positions covered by at least 10 reads from each isolate, with at least 85% of the reads from each isolate supporting a single allele, and with exactly two total alleles over all isolates. SNPs in LTR blocks, as defined above, were removed, leaving 10 SNPs. Alleles ratios were then calculated for each of the 93 non-G217B read alignments for each SNP covered by at least 20 reads for at least 70% of the non-G217B samples, yielding allele ratios for 7 of the 10 original SNPs. At all 7 positions, the majority of reads for the majority of non-G217B samples unanimously assigned a single allele, which was taken as the inferred ancestral G217B allele and a maximum parsimony tree was inferred for UCSF1, UCSF2, and UCSF3, rooted at the inferred G217B. This gave 4 SNPs shared by UCSF1 and UCSF2 on a common branch, 1 SNP unique to UCSF1, assigned to the branch following the divergence of UCSF1 and UCSF2, and 2 SNPs unique to UCSF3, assigned to an independent branch from the root.

### Assessment of mutation rate

Liquid cultures were started from single colonies then passaged once. Cells in this culture were then counted by hemocytometer to calculate total number of cells, which was used to calculate the number of doublings each parental cell had undergone. When cultures were started from single colonies and during the one passage, either all cells from the previous culture or colony were used, or hemocytometer counts were also taken at this intermediate stage to calculate doublings before and after this passage. After parental cultures were grown up, they were sonicated for 2 seconds, diluted, and offspring cells plated for single colonies. Offspring colonies were allowed to grow until very small but clearly visible (about 1 week) at 37 °C then moved to 25 °C and allowed to grow for several weeks. Offspring colonies were then grown up via sequential patching on plates followed by growth in liquid at 37 °C. Offspring genomic DNA was extracted and sequenced as described above. Two parental cultures were also sequenced to confirm our method of inferring parental sequences from offspring. Two previously sequenced strains were ancestral to all parental strains. In total, including data from two experiments, 46 offspring from 11 parental colonies were subjected to whole-genome sequencing for this analysis. There was an average of 31 doublings calculated from parental cells to offspring cells.

Paired-end Illumina reads from the 46 offspring, 2 parental, and 2 ancestral samples were aligned to the UCSF3 reference using BWA MEM 0.7.17 with default parameters. A SNP matrix was defined from the read alignments as nucleotide positions covered by at least 10 reads from each isolate, with at least 85% of the reads from each isolate supporting a single allele, and with exactly two total alleles over all isolates. SNPs in LTR blocks, as defined above, were removed. Additionally, SNPs in chromosome 7, which is often duplicated, were removed so as to consider only haploid positions. The known relationships of the offspring relative to the sequenced and unsequenced parental strains were modeled as a tree, with branches labeled with number of doublings where available (Fig. 3). SNPs were then assigned to branches as follows: if all children of a given parent shared a SNP, then the SNP was assumed to arise at or before the parent; if exactly two children of a given parent differed by a SNP, then the SNP was assigned to the branch of the child that did not match the parent (1 case); if a child differed from all of its siblings by a SNP, then the SNP was assigned to the branch of that child (6 cases); finally, there was one case where the same SNP was observed in two out of 12 siblings. We tried handling this final case in three ways, all of which led to mutation rate estimates that were indistinguishable relative to their 95% confidence intervals: A) the two identical SNPs were each included in calculations, assigned to the branches of both children; B) the SNP was assigned to one child’s branch with the second child’s branch dropped from the calculation; C) the SNP and the branches of both children were dropped from the calculation. These treatments gave a total of 9, 8, or 7 SNPs assigned to branches with measured doublings. The mutation rate was then calculated as the total number of assigned SNPs over the total doublings over all branches, and 95% confidence interval was calculated from the binomial distribution using the binomtest function from SciPy stats^36^. For the above treatments, this gave: A) 2.5873e-10 (95% CI 1.1834e-10 to 4.9122e-10), B) 2.3473e-10 (95% CI 1.0117e-10 to 4.6283e-10), or C) 2.0971e-10 (95% CI 8.4353e-11 to 4.3167e-10).

### Morphology time course conditions and sampling

To obtain yeast and hyphal cells for time courses, 2-day cultures started from single yeast colonies were plated and grown for three weeks both at 25 °C on plates with GlcNAc (which increases the speed and uniformity of hyphal transition^37^) to obtain hyphae and at 37 °C on plates without GlcNAc to obtain paired yeast cells. Prior to the time course, hyphal cultures were started from these plates and grown for 2 days in GlcNAc at 25 °C to ensure full hyphal growth. Subsequently, cultures were passaged 1:6 using media without GlcNAc at 25 °C for 5 days. Cells were confirmed to be fully hyphal by DIC microscopy, and passaged 1:25 and grown for an additional 7 days in media without GlcNAc to ensure steady state hyphal growth. Yeast cells also underwent 1:25 passages once every 2-3 days in liquid media prior to the start of the time course in media without GlcNAc as is standard for *Histoplasma* yeast cultures.

Throughout time courses, all cells were grown in liquid HMM media without GlcNAc, in orbital shakers at 120-150 rpm. Yeast cultures were passaged 1:25 three times a week. Hyphal cultures and morphology transitions were passaged 1:25 once a week. Yeast cultures and cultures undergoing the hyphae-to-yeast transition were grown under standard yeast growth conditions, 37 °C with 5 % CO^2^. Hyphal cultures and cultures undergoing the yeast-to-hyphae transition were grown under standard hyphal conditions, 25 °C with no added CO^2^. Transition cultures were moved to the appropriate conditions on day 0.

DNA sequencing timepoints were collected weekly while steady-state cultures were sampled every two weeks. DNA extraction and sequencing was performed as described above. For microscopy timepoints (taken once a week for the morphology transitions and at the start and end for steady-state growth) cells were fixed in 4 % PFA for 30 minutes at ambient temperature then stored at 4 °C. DIC microscopy images were taken as previously described^34^. 353 microscopy images were randomized then subjected to categorization using the key in Figure S3. At least 4 images were assessed for morphology per transition timepoint and at least 3 were assessed per steady-state timepoint. To ensure consistency, 25 randomly chosen images were shown twice, and each of these images was categorized identically, verifying the analysis pipeline.

### Analysis of variable copy number regions from short read sequencing

Reads were aligned separately to either reference genome UCSF3 or a single copy of the full length LTR transposon sequence used in our published pipelines ^16,18,23,35^. We defined a chromosomal normalizer region for genome UCSF3 as the region between *SRE1* and *MET1* (Chromosome 3 2533970 to 3746857). For each sample, median coverage of the variable coverage region over median coverage in the normalizer region was assessed. The mean and standard deviation of these ratios amongst replicates was plotted.

## Data Availability

Sequencing reads and annotated genome assemblies were submitted to GenBank under BioProjects PRJNA1257453, PRJNA1257851, PRJNA1256922, PRJNA1196486, and PRJNA1196489. Previously published sequencing of natural isolates from BioProjects PRJNA416769, PRJNA868688, and PRJNA1003095 was also used for analysis. Genome assemblies from BioProjects PRJNA682643, PRJNA682644, PRJNA682645, PRJNA682647, and PRJNA682648 were used for comparison. A table listing strains used in this study and a table listing gene names used among Histoplasma assemblies are provided as supplemental tables. Code for SCAR analysis is also provided as a supplemental file.

## Acknowledgements

We would like to thank members of the Sil and Noble labs for helpful discussions. We thank Alexander Johnson, Carol Gross, Joseph Bondy-Denomy, and Suzanne Noble for useful suggestions. We thank Keith Walcott for the sequencing of two strains published here. The UCSF Center for Advanced Technology and the Chan Zuckerburg Biohub-San Francisco supported this research through use of equipment and sequencing.

